# Germline mutagenesis of *Nasonia vitripennis* through ovarian delivery of CRISPR-Cas9 ribonucleoprotein

**DOI:** 10.1101/2020.05.10.087494

**Authors:** Duverney Chaverra-Rodriguez, Elena Dalla Benetta, Chan C. Heu, Jason L. Rasgon, Patrick M. Ferree, Omar S. Akbari

## Abstract

CRISPR/Cas9 gene editing is a powerful technology to study the genetics of rising model organisms, such as the jewel wasp *Nasonia vitripennis*. However, current methods involving embryonic microinjection of CRISPR reagents are challenging. Delivery of Cas9 ribonucleoprotein into female ovaries is an alternative that has only been explored in a small handful of insects, such as mosquitoes and whiteflies. Here, we developed a simple protocol for germline gene editing by injecting Cas9 ribonucleoprotein in adult *N. vitripennis* females using either ReMOT control (Receptor-Mediated Ovary Transduction of Cargo) or BAPC (Branched Amphiphilic Peptide Capsules) as ovary delivery methods. We demonstrate efficient delivery of protein cargo such as EGFP and Cas9 into developing oocytes via P2C peptide and BAPC. Additionally, somatic and germline gene editing have been demonstrated. This approach will greatly facilitate CRISPR-applied genetic manipulation in this and other rising model organisms.

## Introduction

Hymenoptera is an extremely prominent insect order that includes several thousand species of ants, bees and wasps. The jewel wasp, *Nasonia vitripennis*, is one of the most widely studied hymenopteran species. It is one of four species in the *Nasonia* genus, all of which parasitize the pupae of the blowfly *Sarcophaga bullata. N. vitripennis* is easily maintained in the laboratory environment due to its short generation time (^~^2 weeks at 25°C) and can be easily reared in small vials with *S. bullata* pupae (Werren and Loehlin, 2009). Since 1980 *N. vitripennis* has been used as a comparative model insect for the study of sex ratio control (Parker and Orzack, 1985; Werren, 1983), sex determination (Beukeboom and Kamping, 2006; Beukeboom et al., 2007), speciation (Breeuwer and Werren, 1995; Buellesbach et al., 2013; Ellison et al., 2008) and host-parasite evolution (Bordenstein et al., 2001; Breeuwer and Werren, 1990). *Nasonia* is particularly amenable for genetic studies due to advantages provided by its haplodiploid sex determination system, such as the easy screening of recessive mutations in haploid males (Werren and Loehlin, 2009). Recently, *Nasonia* also has been used as a model organism for addressing questions related to the selfish B chromosome function (Akbari et al., 2013; Aldrich et al., 2017; Dalla Benetta et al., 2020; Ferree et al., 2015), polyploidy (Leung et al., 2019), circadian and photoperiodic response (Dalla Benetta et al., 2019; Paolucci et al., 2019), and genomic imprinting (van de Zande and Verhulst, 2014; Verhulst et al., 2013, 2010).

Many molecular tools are available for the study of phenomena in *N. vitripennis;* these include fluorescent *in situ* hybridization (FISH) (Larracuente and Ferree, 2015) and transient, systemic RNA interference (RNAi) (Lynch and Desplan, 2006). However, perhaps the most promising tool, CRISPR/Cas9 gene editing, has recently been developed in *N. vitripennis* (Li et al., 2017a). Specifically, CRISPR was used to induce site specific mutations in several visible marker genes, demonstrating the promise of this method for gene editing in this organism. However, current techniques for the application of CRISPR in *N. vitripennis* require microinjection into preblastoderm embryos in order to generate heritable mutations, thus requiring specialized equipment and skills (Li et al., 2017b). *Nasonia* eggs are very small (0.1-0.16 mm) and require transplantation of injected embryos into a pre-parasitized blowfly pupa for successful development (Li et al., 2017b). In addition to the small egg size and requirement for transplantation, the presence of a viscous cytoplasm causes frequent clogging of the needle (Li et al., 2017b), making *Nasonia* egg microinjection even more challenging. The difficulties of efficiently injecting embryos, combined with the difficulties of follow-up screening of mutants in case of target genes that do not present visible phenotypes, dramatically limits the use of CRISPR in *Nasonia* to highly specialized research groups. Thus, there is a strong need for easier CRISPR reagent delivery approaches in these species.

Recently, two alternative methods that bypass the requirement for embryonic microinjection have been developed (Chaverra-Rodriguez et al., 2018; Dermauw et al., 2020; Heu et al., 2020; Hunter et al., 2018; Macias et al., 2020). Receptor-Mediated Ovary Transduction of Cargo (ReMOT)involves delivery of the CRISPR/Cas9 ribonucleoprotein (RNP) complex (Cas9 with a single-guide RNA (sgRNA)) into insect ovaries. This approach has been successfully used to create heritable gene edits in the germline of several mosquito species (Chaverra-Rodriguez et al., 2018; Macias et al., 2020) and also recently in the silverleaf whitefly (Heu et al., 2020). This approach employs the use of peptide ligands derived from yolk protein precursors (YPPs) fused to the Cas9-RNP complex. During vitellogenesis, the process of ovary and egg maturation(Valle, 1993), YPPs become synthesized in the fat body, secreted into the hemolymph, and delivered into the ovaries by receptor-mediated endocytosis (Raikhel and Dhadialla, 1992). Therefore, injection of the ReMOT-RNP complex into the hemolymph of vitellogenic females enabled efficient receptor-mediated delivery of the CRISPR machinery into developing oocytes, resulting in targeted gene editing in embryos, thus bypassing the need for microinjection (Chaverra-Rodriguez et al., 2018).

An alternative approach, termed BAPC-assisted CRISPR delivery, involves the use of Branched Amphiphilic Peptide Capsules BAPC-tofect™ (Phoreus™ Biotechnology, Inc.) for delivery of CRISPR ribonucleoprotein into the ovary. These peptide nanospheres consist of equimolar proportions of two branched peptide sequences, bis(FLIVI)-K-KKKK and bis(FLIVIGSII)-KKKKK, which when added together, self-assemble to form bi-layer capsules (Sukthankar et al., 2014). The capsules are water soluble, stable in blood, and made entirely of natural amino acids (Sukthankar et al., 2014). The size of the capsules range from 10–50 nm (depending on annealing) and are resistant to detergents and proteases (Avila et al., 2016; Sukthankar et al., 2013). Additionally, the capsules interact with DNA and RNA acting as cationic nucleation centers with the negatively charged DNA binding to the outer surface. Physical BAPC/DNA interactions result in compact clusters ranging in average size from 50 to 250 nm (Avila et al., 2016; Sukthankar et al., 2013). BAPC can thus facilitate the uptake of DNA and potentially other types of nucleic acid *in vitro* and *in vivo*. It has been used for delivery of dsRNA and plasmid DNA in cultured insect and animal cells (Avila et al., 2018; Barros et al., 2016) and recently it has been used to improve the delivery of CRISPR components into adult ovaries to facilitate heritable gene editing in the Asian citrus psyllid *Diaphorina citri (Hunter et al., 2018)*.

These new methods, involving the delivery of CRISPR reagents into adults that facilitate ovary uptake, can simplify and possibly enable broader use of CRISPR technology in the jewel wasp *N. vitripennis* and other hymenoptera. Here we report the optimization of ReMOT- and BAPC-mediated CRISPR/Cas9 in *N. vitripennis*. We demonstrate efficient delivery of Cas9 RNP to *N. vitripennis* ovaries by either P2C peptide (Chaverra-Rodriguez et al., 2018) or BAPC capsules, each resulting in both somatic and heritable germline-induced mutations without requiring embryonic microinjections.

## Results

### Delivery of P2C-EGFP-Cas9 into Nasonia ovaries

To determine if P2C could deliver EGFP protein efficiently into ovaries, we injected 30 black *Nasonia vitripennis* pupae, one day pre-emergence, with P2C-EGFP and 30 with EGFP without P2C. We observed efficient protein delivery by P2C in ovaries 24 and 48h post injection (p.i.) and in embryos laid 72h post injection (p.i.) (Fig. 1; Fig. S1, S2). Although fluorescence was also detected in P2C-EGFP injected ovaries both at 24 and 48h p.i. (Fig. 1A), we only counted positive EGFP oocytes at 48 and 72h p.i. because we noted that injected wasps usually do not lay eggs within the 24h post injection. At 48h p.i., fluorescence was detected in 29 oocytes across 10 ovaries (Fig. 1A, S1A). The fluorescence was localized at the terminal portion of the ovarioles (Fig. 1A), and not detected in all oocytes from the same ovary. EGFP was not detected at 72h p.i. (Fig. 1A, Fig. S1A). Ovaries injected with EGFP without the P2C ligand showed some background fluorescence, but it was not localized as P2C-EGFP (Figure 1A, S1B). We also screened fluorescence in embryos laid 72h p.i. (Fig. 1A, Fig. S2). About 50% (6 out of 12) of P2C-EGFP embryos showed EGFP presence indicating an efficient delivery of protein from P2C peptide into *Nasonia* eggs. Interestingly, we detected some low background fluorescence in about 15% (2 out of 13) of embryos, from the group injected only with EGFP (without P2C), (Fig. 1A, Fig. S2). No fluorescence was detected in ovaries and embryos of non-injected wasps (Fig. 1A, S 1C, S2).

**Figure 1.**
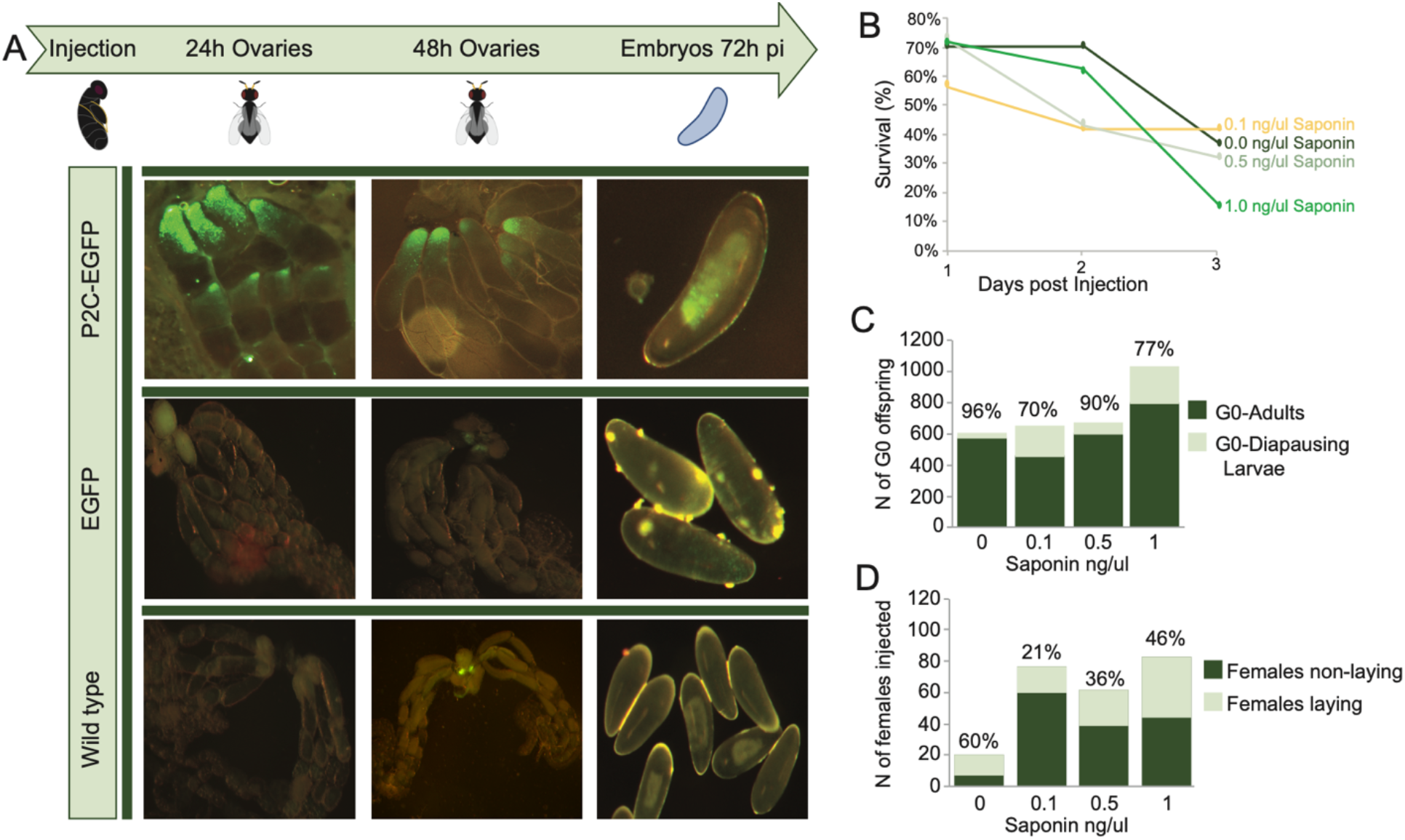
Injection optimization. **A)** Delivery of EGFP by the peptide P2C into the ovaries of *N. vitripennis* at 24, 48 hours post injection (p.i.) and in embryos laid 72h p.i. **B)** Effect of saponin on survival of injected females. **C)** Effect of saponin on G0 offspring. **D)** Effect of saponin on number of females laying eggs.

### Optimization of Saponin injection

Saponin was successfully used as an endosomal escape reagent, in order to facilitate the release of RNP into the ovaries in mosquitoes (Chaverra-Rodriguez et al., 2018, Macias et al., 2020). However it can have detrimental effects on the fitness of the injected individuals (Heu et al., 2020). Therefore, preliminary injections at three different concentrations of saponin (0.1, 0.5 and 1 mg/ml) were carried out to test its effects on female survival and fecundity (Fig. 1B-D, Table S1). No differences were detected on survival across three days among treatments. Between 58 to 72% of the females emerged 24h p.i. and on day 3 p.i between 21 to 45% were still alive (Fig. 1B, Table S1). However, a low percentage of injected females laid eggs (21 to 60%), without a direct correlation with saponin concentration (fig. 1C, Table S1). There was no clear effect of saponin on G0 offspring survival. However, at the concentration of 1 mg/ml only 46% of G0 individuals developed to adulthood, whereas the rest stopped their development at the larval phase (Fig. 1D, Table S1).

### Somatic and germline gene editing of *cinnabar* gene

In order to determine whether ReMOT or BAPC are capable of facilitating CRISPR gene editing through delivery of CRISPR ribonucleoproteins into developing eggs, we injected black pupae (1 day prior emergence) and 1 to 2 day-old adult females with varying concentrations of Ribonucleoprotein (RNP): P2C-EGFP-Cas9, P2C-Cas9-sgRNA or BAPC-Cas9 combined with sgRNA targeting *NVcin*. The target gene, *NVcin*, encodes kynurenine hydroxylase, an enzyme involved in ommochrome biosynthesis, which is required for dark eye pigmentation (Sethuraman and O’brochta, 2005). Cas9-sgRNA (without delivery reagent) and non-injected wasps were used as negative controls. Non-injected wasps produced only wild type offspring, whereas 35 out of 65 Cas9-sgRNA injected wasps produced one somatic mosaic mutant and one off-target germline mutant out of 2690 offspring (Fig. S4, Table 1, Table S2). When we injected RNP P2C-EGFP-Cas9 at concentrations between 400-500 ng/ul and saponin at 0.1-0.5 mg/ml in virgin females, we observed 34 females laying eggs out of 123 females injected. Among those 34 females, 7 females laid 8 mutant offspring with mild phenotypes for *cinnabar* regardless of the female age at the time of injection (Table 1, Figure 2A). *NVcin* sequences from these mutants were mostly wild type, with only one individual that showed a secondary peak in the target area of the sgRNA indicating the mosaic nature of those mutants (Fig. 2B). However, increasing the concentration of P2C-Cas9 protein/sgRNA to 3000 ng/ul (see Table S2 for concentration details) resulted in 45 out of 121 injected individuals that laid eggs. Four of these 45 females produced 3 full *cin* mutants with bright red eyes (Table 1, Fig. 2A) and one with the mild, red-eye phenotype. Subsequent sequencing of *NVcin* with the bright red eyes confirmed efficient gene editing of the target gene (Fig. 2B). These data indicated that higher concentration of what RNP and absence of EGFP might help increase gene editing events. Injection of BAPC-RNP with different concentrations of protein/sgRNA ranging from 360 to 500 ng/ul was also efficient in generating *cin* mutants either with mild or strong red phenotype (Fig 2A, Table 1, Table S2). 32 out of 60 injected females efficiently laid eggs and 5 females produced 7 mutants. However only one individual showed heritable gene editing. (Fig 2B, Table 1).

**Figure 2.**
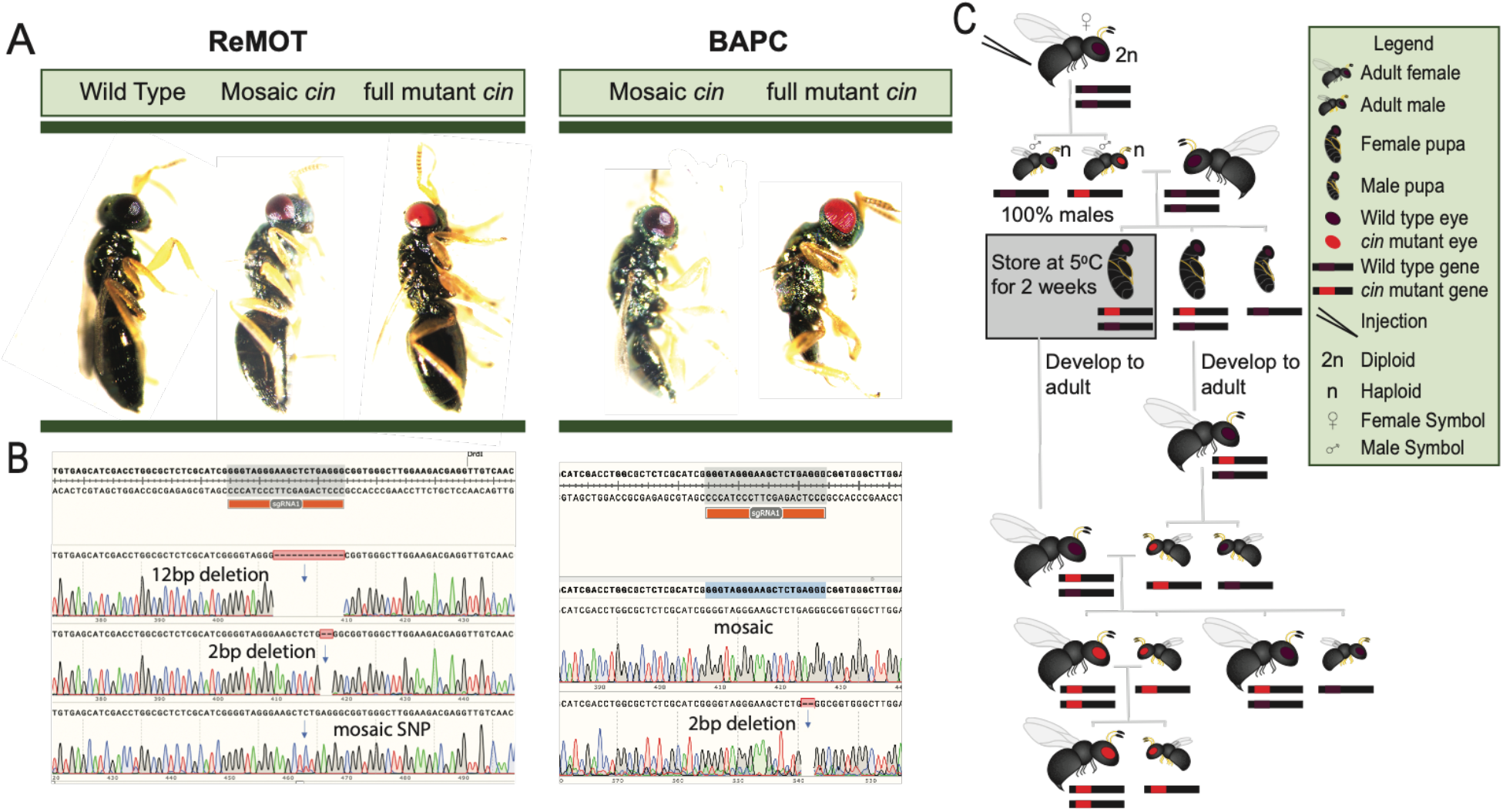
Gene editing mediated by ReMOT and BAPC. **A)** Diversity of phenotypes observed in G0 males from virgin females injected either with ReMOT Control or BAPC. Wild type is the representative result of the two negative controls (non-injected and CAS9-sgRNA-injected wasps) **B)** Sequencing results of the target region of the *NVcin* locus of G0 mutant males. **C)** Crossing scheme to distinguish somatic from germline mutants and stabilize a mutant colony with red eyes.

**Table 1.**
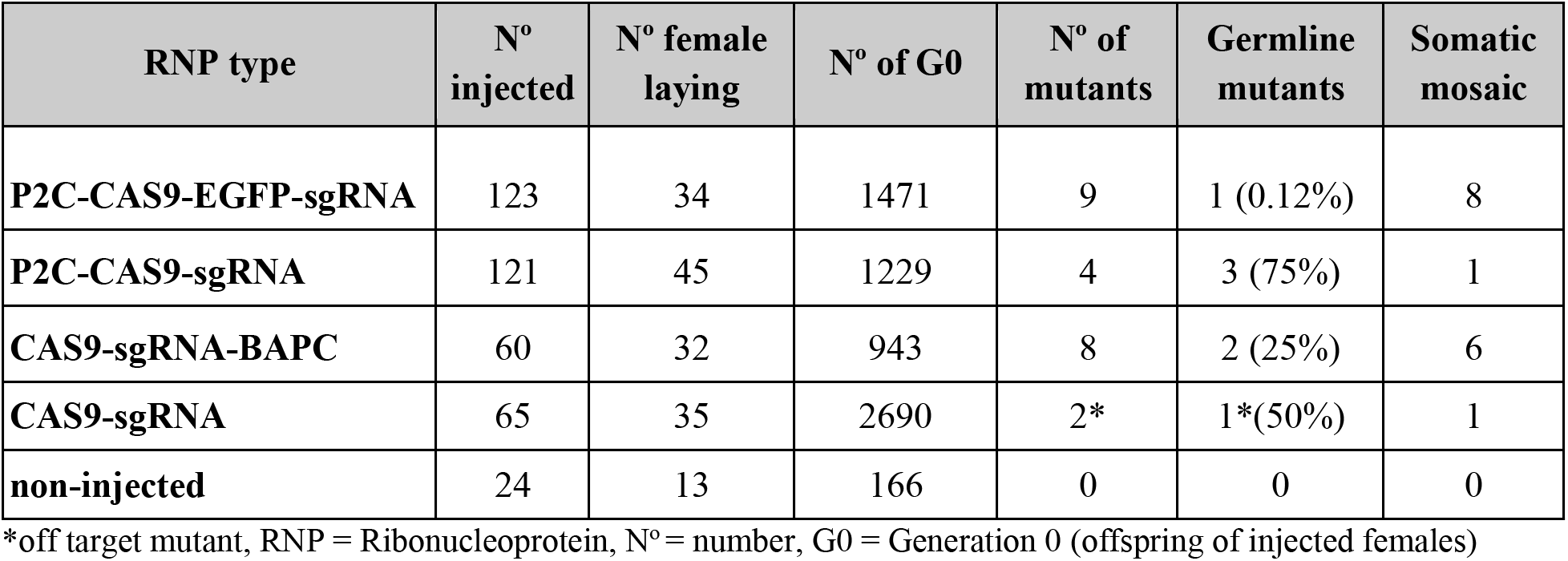
Experiments testing gene editing in *Nasonia vitripennis*

| RNP type | N° injected | N° female laying | N° of G0 | N° of mutants | Germline mutants | Somatic mosaic |
| --- | --- | --- | --- | --- | --- | --- |
| **P2C-CAS9-EGFP-sgRNA** | 123 | 34 | 1471 | 9 | 1 (0.12%) | 8 |
| **P2C-CAS9-sgRNA** | 121 | 45 | 1229 | 4 | 3 (75%) | 1 |
| **CAS9-sgRNA-BAPC** | 60 | 32 | 943 | 8 | 2 (25%) | 6 |
| **CAS9-sgRNA** | 65 | 35 | 2690 | 2* | 1*(50%) | 1 |
| **non-injected** | 24 | 13 | 166 | 0 | 0 | 0 |

\*off target mutant, RNP = Ribonucleoprotein, N° = number, G0 = Generation 0 (offspring of injected females)

Germline mutants were distinguished from somatic mutants by subsequent crosses of *cin* males to wild type females (Fig. 2C). The G1 heterozygotes females produced by these crosses had wild type eyes and were allowed to lay unfertilized eggs (Fig 2C). Unfertilized eggs in *Nasonia* lead to haploid males, allowing the detection of mutants. G2 male progeny with the *cin* phenotype were generated only from G0 males with bright red eyes (Fig. 2C, Table 1), whereas no mild phenotypes were detected in G2 offspring. These data confirm that the mild color phenotype was due to somatic mosaicism while the bright red eye phenotype resulted from gene editing in the germ line.

## Discussion

ReMOT control (Receptor-Mediated Ovary Transduction of Cargo) was recently developed as an effective means for mediating CRISPR/Cas9 gene editing in mosquitoes (Chaverra-Rodriguez et al., 2018)(Chaverra-Rodriguez et al., 2018; Dermauw et al., 2020; Heu et al., 2020; Hunter et al., 2018; Macias et al., 2020)(Chaverra-Rodriguez et al., 2018) and whiteflies (Heu et al., 2020). The approach was to use a yolk protein peptide fused to a molecular cargo in order to deliver the CRISPR ribonucleoprotein into female ovaries. Here, we demonstrate the efficiency of the *Drosophila melanogaster* derived P2C peptide for ovary uptake in *Nasonia vitripennis*, allowing us to bypass the challenges of embryonic microinjections to induce heritable genome editing. The P2C peptide was initially optimized for protein cargo delivery in Aedes (Chaverra-Rodriguez et al., 2018) and *Anopheles* (Macias et al., 2020) mosquito ovaries, but was unsuccessful in the silverleaf whitefly, and therefore a native yolk protein-derived peptide (BtKV) was produced for efficient delivery (Heu et al., 2020). Interestingly, we have demonstrated that the protein EGFP lacking the P2C peptide was transduced into *N. vitripennis* ovaries, although the detection was very low in frequency and signal intensity. Similarly, the injection of Cas9 RNP without P2C resulted in gene editing at a lower rate compared to the proteins P2C-EGFP-Cas9 or P2C-Cas9. This pattern was previously reported for mosquitoes (Chaverra-Rodriguez et al., 2018) showing that vitellogenic ovaries of two different insect orders may uptake proteins from the hemolymph by non-specific endocytosis. Here, we demonstrate that both proteins, EGFP and Cas9 fused to P2C were delivered more efficiently into ovaries and consequently into embryos for at least 72h p.i. This result, and a recent report of P2C mediated delivery of the protein mCherry into the ovaries of the asian citrus psyllid *Diaphorina citri* (Chaverra-Rodriguez et al., 2020) suggest that P2C might be effective for effective delivery of ribonucleoprotein into the oocytes of a wide range of different insect species.

We have demonstrated that ReMOT control can be used in *N. vitripennis* for targeted and heritable gene editing, without requiring embryonic microinjections. We used two different P2C-fusion proteins. The use of high ribonucleoprotein protein (RNP, P2C-Cas9-sgRNA) concentration of about 3000 ng/ul, was able to induce germline gene editing, whereas lower RNP concentration gave rise mainly to somatic mosaicism. Additionally, the fusion of EGFP on P2C-Cas9 reduced the efficiency of germline mutation. Lower gene editing efficiency from P2C-EGFP-Cas9 compared to P2C-Cas9 was also described in mosquitoes (Chaverra-Rodriguez et al., 2018) and it was hypothesized that it could be a consequence of steric interactions between the proteins. Using this approach, we were able to generate somatic and germline *cinnabar* mutations up to 3 days p.i., indicating that the CRISPR system is active both before and after embryogenesis, as has been demonstrated in the other insects (Chaverra-Rodriguez et al., 2018). Further optimization of ReMOT technology in this species could be achieved by producing fusion proteins of Cas9 and the native yolk proteins of *N. vitripennis* acting as ligands for ovary receptors, or by modifying the concentrations of protein and endosomal escape reagents. In mosquitoes, an endosomal escape reagent like chloroquine or saponin is necessary to achieve efficient gene editing (Chaverra-Rodriguez et al., 2018; Macias et al., 2020). Endosomal escape reagents help the release of the ribonucleoprotein once delivered into the ovary (Fuchs et al., 2013). However, in the whitefly the use of saponin was unnecessary, and even detrimental for viability (Heu et al., 2020). Here, we used different concentrations of saponin to mediate endosomal escape and, aside from some mild effect on survival and fecundity, its use did not seem to be detrimental in *Nasonia*. However, high concentrations of saponin increased the production of diapause larvae. Diapause response in *Nasonia* is maternally induced and often linked to stress response (Saunders, 1965), indicating a possible toxic effect of saponin at higher concentrations. Moreover, unlike in mosquitoes, a very low concentration (0.1 ng/ul) was sufficient for gene editing, indicating that the use of endosomal escape needs to be optimized for each species.

The second method we used to achieve targeted gene editing involved the use of nanosphere peptides that self-assembled to form capsules able to bind and deliver ribonucleotides and RNP across membranes. Specifically, we used BAPC (Branched Amphiphilic Peptide Capsules BAPC-tofect) to encapsulate the ribonucleoprotein Cas9-sgRNA. We demonstrate that BAPC was able to deliver RNP into the ovary and induce targeted gene editing both before and after embryogenesis like ReMOT Control-mediated CRISPR/Cas9 delivery. Contrary to ReMOT Control, BAPC required a lower concentration of RNP ranging from 360 to 500 ng/ul, comparable to the one used for embryonic microinjection (Li et al., 2017b), therefore suggesting that higher concentrations of BAPC-RNP may increase gene editing efficiency. In addition, no endosomal escape is required to induce editing events, although its use in conjunction with BAPC has not been tested. These data suggest that BAPC is an effective alternative to ReMOT Control for insect species in which P2C peptide is not able to deliver protein cargo or in which the yolk peptides are not known or difficult to produce or obtain. Importantly BAPC is also able to bind different types of ribonucleotides like plasmid DNA and dsRNA, acting as cationic nucleation centers with the negatively charged DNA binding to the outer surface (Avila et al., 2018, 2016; Barros et al., 2016). This opens up the possibility of combining RNP with template DNA to promote transgenesis mediated by homology-directed repair (HDR) events induced by CRISPR/Cas9, thus circumventing embryonic microinjection. In *N. vitripennis*, transgenesis still needs to be demonstrated, and embryonic microinjection remains as a formidable obstacle (Li et al., 2017b). Therefore, BAPC may allow for the development of transgenic strategies, either through CRISPR/Cas9 HDR-mediated or transposon-mediated (Kimura, 2001), in order to express transgenes and increase functional studies in this species.

Taken together we have demonstrated gene editing in *Nasonia* that is mediated by two separate approaches that involve injecting adults as alternatives to embryonic microinjection. This opens the possibility of increasing the use of CRISPR/Cas9 in *N. vitripennis* and other insect species for which embryonic microinjection is challenging. For example, several *D. melanogaster* parasitoids, which belong to the genus *Leptopilina*, deposit single eggs into a single *Drosophila* larva. Eggs are therefore impossible to reach and inject in those species. Additionally, several optimizations of the system in those emerging model systems, like adjusting the concentration of reagents, increasing the number of injected wasps and limiting the time of egg laying will help to identify mutants more easily and therefore be applied to functional studies of genes with not visible phenotypes.

### Experimental procedures

#### *Nasonia vitripennis* strain and rearing

Experiments were performed using the lab strain AsymCX, which genome has been sequenced (Dalla Benetta et al., 2020). Wasps were maintained on *Sarcophaga bullata* pupae as hosts in mass culture vials under LD 12:12, 26 ± 1°C. In order to test what was the best time to deliver protein into the ovaries twelve days black pupae (one day prior to emergence) and one to two days adult females were used for our experiments. Black pupae were maintained at 4°C until injections in order to stop development. Gene editing was tested in virgin females because unfertilized eggs give rise to haploid male and allow an easy screening of the recessive phenotype.

#### Injections to detect ovary intake of fluorescent proteins by P2C

To detect if the previously used P2C peptide (Chaverra-Rodriguez et al., 2018) is efficient to deliver proteins into *Nasonia* ovaries, we injected 30 female pupae with P2C-EGFP and 30 female pupae with EGFP (without the P2C ligand). In addition, 30 non-injected female pupae were used as negative control. To visually confirm EGFP in the ovaries we performed dissections at 24, 48- and 72-hours post injection (p.i.) of 10 pairs of ovaries per treatment. Dissected ovaries from 48 and 72 hours were mounted in a saline buffer mixed with SlowFade Gold^®^ antifade reagent (Invitrogen), covered with a coverslip and imaged on an Olympus BX41 epifluorescent microscope. At 48 and 72h p.i., we counted the number of oocytes with fluorescence signal in each ovary. Additionally, embryos produced from females 72h p.i. were screened for the presence of EGFP.

#### Protein preparation

The plasmids pET28a-P2C-EGFP (Chaverra-Rodriguez et al., 2018), pET28a-EGFP (Chaverra-Rodriguez et al., 2018), pET28a-P2C-Cas9 (Chaverra-Rodriguez et al., 2018) and pET28a-P2C-EGFP-Cas9 (Chaverra-Rodriguez et al., 2018) were transformed into *E. coli* BL21 (NEB) and transformation was verified by PCR and sequencing. To induce expression of recombinant protein, a preculture was grown overnight at 37°C in 50 ml of LB medium supplemented with kanamycin (100 μg/μl). After 12 hours, 10 ml of preculture was transferred to 990 ml of LB supplemented with the same antibiotics, incubated at 37°C until OD=0.6, when IPTG was added at a final concentration of 0.5 mM, the culture was incubated for 20 h at 18°C. Cells were spun down at >10,000 xg for 15 min, resuspended in 50 ml lysis buffer (20 mM Tris-HCl pH 8.0, 300 mM NaCl, 10 mM imidazole), placed at −80°C overnight and it was sonicated 5 times at 60% duty 5 sec pulse 5 sec rest (two aliquots each 25 ml), centrifuged at 13,000 xg for 30 min, the supernatant removed and incubated with Ni-NTA beads with rotation at 4°C for 2 hours. Beads were washed 3 times with 10 ml wash buffer (20 mM Tris-HCl pH 8.0, 300 mM NaCl, 30 mM imidazole) and eluted with 1 ml elution buffer (20 mM Tris-HCl pH 8.0, 300 mM NaCl, 250 mM imidazole) 10 times. Eluted protein was dialyzed in a Slide-A-Lyzer dialysis cassette (Thermo Fisher) for two hours in dialysis buffer (50 mM Tris-HCl pH 8.0, 300 mM KCl, 0.1 mM EDTA, 0.5 mM PMSF) and buffer changed every 2 hours 2 times, then left overnight at 4°C with gentle agitation in fresh buffer. Protein purity was visualized by SDS-PAGE.

#### *In vitro* cleavage assay

To test the cleavage activity of the produced P2C-Cas9 before injection we performed an *in vitro* cleavage assay. First, we extracted the DNA from an adult wasp using DNeasy Blood & Tissue Kit (Qiagen #69504) following the manufacturing protocol. We then performed a PCR of the target gene *cinnabar* using forward primer 5’-TTTCGCTATACTCTCTCGTGTG-3’ and reverse primer 5’-GAGCGAGACTCGAGCAATAAC-3’ and Q5 polymerase (New England Biolabs, #M0491S). PCR was run using the following conditions: 30 sec at 98° C followed by 35 cycles of 10 sec at 98° C, 30 sec at 58° C and 60 sec at 72° C, followed by 5 min at 72° C. To test the activity of P2C-Cas9; 300 ng of P2C-Cas9 and 300 ng of sgRNAs were combined with 70 ng of purified PCR fragment and were diluted in 1 μL of NEB buffer 3.1 to a total volume of 10 μL in water. Reactions were run for 1h at 37°C and diagnostic bands visualized by electrophoresis in a 1% agarose/TAE gel. The protein batches that showed efficient cleavage were then used in the following steps (Fig. S1).

#### Females injection mix preparation for ReMOT gene editing

To find the most efficient amount of protein/sgRNA to induce germline gene editing, different concentrations of ribonucleoprotein (either Cas9, P2C-Cas9-EGFP or P2C-Cas9 combined with sgRNA) were tested. We used the sgRNA targeting *cinnabar* (NV14284) that was efficiently mutated using embryonic microinjection CRISPR (Li et al., 2017a). The sgRNA was synthesized by Synthego and was prepared in aliquots of 500 ng/ul and 5000 ng/ul and stored at −80° C. The protein (CAS9, P2C-Cas9-EGFP or P2C-Cas9) was mixed with sgRNA at different concentrations and volumes considering that the molar ratio sgRNA to protein was >2 and the final concentration of protein was between 500 to 3000 ng/μl. After mixing the RNP was incubated at room temperature for 15 min and then placed on ice. Immediately before the injection the endosomal escape reagent saponin was added at a final concentration of 0.1-1 mg/ml. The mix was mixed slowly and used immediately for injections. In order to identify the highest concentration of saponin that allows good survival and fecundity of injected individuals, we perform a series of preliminary injections using different concentrations of saponin. 40 pupae were injected respectively with 0.1, 0.5 or 1 mg/ml of saponin and survival was recorded for 3 days post injection. At 24h p.i., surviving females emerged from their puparium and were allowed to oviposit on one *S. bullata* pupae for 24h and then re-hosted 48-72 hours later. Number of G0 offspring that developed to adult as well as diapausing larvae were counted per each host. Diapause in *Nasonia* is maternally induced and also used as a stress response(Saunders, 2014). Higher number of diapausing larvae could thus indicate a toxic effect of saponin. Additionally, the percentage of female laying eggs can also be used as a quantification for saponin toxicity. Saponin from Quillaja saponaria molina (Acros Organics, C41923-1000) stocks were prepared freshly prior to each injection by diluting 10 mg of saponin in 200 ul of PBS 1X. This stock solution was used to prepare fresh dilutions at the concentrations required.

#### Females injection mix preparation for BAPC gene editing

Branched Amphiphilic Peptide Capsules BAPCtofect™ was purchased from Phoreus™ Biotechnology, Inc. (#B25005, 0.5 mg). 10 μg/μL stock solutions were prepared by rehydrating BAPCtofect-25 with 50 μL of nuclease free water, and rehydrating CaCl2 with 100 μL of nuclease free water. Cas9 (PNABio #CP01-20) and sgRNA were mixed at different concentrations and volumes resulting in a final concentration per each component ranging between 360-600 ng/μl. The mixture was then incubated at room temperature for 15 min to allow the formation of the RNP complex and then placed on ice for 10 min. BAPC was then added to reach a final concentration of 20 ng/μl and incubated at room temperature for 30 min. Finally, 1 μl of CaCl2 was added to the solution and incubated for another 10-20 min prior to injection. Solution was then kept in ice and injected. Control treatments include Cas9-sgRNA (360 ng/μl each) without BAPC and noninjected wasps.

#### Female microinjections

Adults and pupae female injections were performed with an aspirator tube assembly (A5177, Sigma) fitted with a glass capillary needle. Pupae were aligned and attached to a glass slide using glue. Adult females were immobilized at 4°C following and kept on ice during injection. Females and pupae were injected in the abdomen until no additional fluid would enter, at an approximate volume of 200 nl per female.

#### Screening and maintenance of mutants

Injected females were singularly reared on a *S. bullata* pupae for 24 hours and then re-hosted 48-72 hours later. The G0 progenies, resulting in 100% male broods (haplodiploid sex determination), were reared to adults and screened for eye color changes using a dissecting microscope. The individuals that showed red color in the eyes were allowed to mate to wildtype females, the G1 females coming from this outcross were allowed to lay unfertilized G2 progeny (haplodiploidy and thus only male offspring) that were screened again for eye color. A few of G1 females were stored at 5° C in the pupal stage until G2 progeny developed and then crossed with red-eye G2 males to allow the generation of a mutant *cin* colony at G3 (Fig. 2C). Both G0 and G2 individuals were used for molecular characterization (see below).

#### Molecular confirmation of INDELs in the target gene

To confirm the presence of indels on G0 and G2 individuals, DNA extraction was performed with Qiagen kit (DNeasy Blood & Tissue Kit) and PCR was performed using primers and conditions as described above. PCR samples were purified with MinElute PCR Purification Kit (Qiagen #28004) and sent for sequencing to Retrogen. Sequencing results were aligned using Snapgene to verify the presence of INDELs or SNPs.

## Acknowledgements

This work was supported in part by funding from generous UCSD laboratory startup funds awarded to O.S.A. NSF/BIO grant 1645331, NIH/NIAID grant R21AI111175, USDA/NIFA grant 2014-10320, USDA Hatch funds (Accession #1010032; Project #PEN04608), and a grant with the Pennsylvania Department of Health using Tobacco Settlement Funds to J.L.R, and NSF funding awarded to P.M.F. (NSF-1451839).

## Author Contributions

DCR, EDB, PMF and OSA conceived the project and supervised the research. CCH and JLR produced the proteins needed to conduct the experiments. DCR and EDB conducted the experiments. DCR and EDB analyzed the data. All authors contributed to writing and have reviewed and approved of the manuscript prior to submission. The manuscript has been submitted solely to this journal and is not published, in press, or submitted elsewhere.

## Data Availability

All data are described in the text and/or included in tables and figures within the manuscript and in the supplementary information files including:

**Table S1 Saponin injections**

**Table S2 Ribonucleoprotein injections (separate file)**

**Fig. S1 *In vitro* cleavage assay**

**Fig. S2 Ovaries intake**

**Fig. S3 Embryos from 72h p.i. Wasps**

**Fig. S4 Cas9-sgRNA control injections**

## References

Akbari, O.S., Antoshechkin, I., Hay, B.A., Ferree, P.M., 2013. Transcriptome profiling of *Nasonia vitripennis* testis reveals novel transcripts expressed from the selfish B chromosome, paternal sex ratio. G3 3: 1597–1605.

Aldrich, J.C., Leibholz, A., Cheema, M.S., Ausió, J., Ferree, P.M., 2017. A “selfish” B chromosome induces genome elimination by disrupting the histone code in the jewel wasp *Nasonia vitripennis*. Sci. Rep. 7: 42551.

Avila, L.A., Aps, L.R.M.M., Ploscariu, N., Sukthankar, P., Guo, R., Wilkinson, K.E., Games, P., Szoszkiewicz, R., Alves, R.P.S., Diniz, M.O., Fang, Y., Ferreira, L.C.S., Tomich, J.M., 2016. Gene delivery and immunomodulatory effects of plasmid DNA associated with Branched Amphiphilic Peptide Capsules. J. Control. Release 241: 15–24.

Avila, L.A., Chandrasekar, R., Wilkinson, K.E., Balthazor, J., Heerman, M., Bechard, J., Brown, S., Park, Y., Dhar, S., Reeck, G.R., Tomich, J.M., 2018. Delivery of lethal dsRNAs in insect diets by branched amphiphilic peptide capsules. J. Control. Release 273: 139–146.

Barros, S.M., Whitaker, S.K., Sukthankar, P., Avila, L.A., Gudlur, S., Warner, M., Beltrão, E.I.C., Tomich, J.M., 2016. A review of solute encapsulating nanoparticles used as delivery systems with emphasis on branched amphipathic peptide capsules. Arch. Biochem. Biophys. 596: 22–42.

Beukeboom, L.W., Kamping, A., 2006. No patrigenes required for femaleness in the haplodiploid wasp *Nasonia vitripennis*. Genetics 172: 981–989.

Beukeboom, L.W., Kamping, A., van de Zande, L., 2007 Sex determination in the haplodiploid wasp *Nasonia vitripennis* (Hymenoptera: Chalcidoidea): A critical consideration of models and evidence. Semin. Cell Dev. Biol. 18: 371–378.

Bordenstein, S.R., O’Hara, F.P., Werren, J.H., 2001. *Wolbachia*-induced incompatibility precedes other hybrid incompatibilities in *Nasonia*. Nature 409: 707–710.

Breeuwer, J.A.J., Werren, J.H., 1995. Hybrid breakdown between two haplodiploid species: the role of nuclear and cytoplasmic genes. Evolution 49: 705–717.

Breeuwer, J.A., Werren, J.H., 1990. Microorganisms associated with chromosome destruction and reproductive isolation between two insect species. Nature 346: 558–560.

Buellesbach, J., Gadau, J., Beukeboom, L.W., Echinger, F., Raychoudhury, R., Werren, J.H., Schmitt, T., 2013. Cuticular hydrocarbon divergence in the jewel wasp *Nasonia:* evolutionary shifts in chemical communication channels? J. Evol. Biol. 26: 2467–2478.

Chaverra-Rodriguez, D., Macias, V.M., Hughes, G.L., Pujhari, S., Suzuki, Y., Peterson, D.R., Kim, D., McKeand, S., Rasgon, J.L., 2018. Targeted delivery of CRISPR-Cas9 ribonucleoprotein into arthropod ovaries for heritable germline gene editing. Nat. Commun. 9: 3008.

Dalla Benetta, E., Antoshechkin, I., Yang, T., Nguyen, H.Q.M., Ferree, P.M., Akbari, O.S., 2020. Genome elimination mediated by gene expression from a selfish chromosome. Sci Adv 6: eaaz9808.

Dalla Benetta, E., Beukeboom, L.W., van de Zande, L., 2019. Adaptive differences in circadian clock gene expression patterns and photoperiodic diapause induction in *Nasonia vitripennis*. Am. Nat. 193: 881–896.

Dermauw, W., Jonckheere, W., Riga, M., Livadaras, I., Vontas, J., Van Leeuwen, T., 2020. Targeted mutagenesis using CRISPR-Cas9 in the chelicerate herbivore Tetranychus urticae. Insect Biochem. Mol. Biol. 120: 103347.

Chaverra-Rodriguez, D., Bui, M., Li, M., Raban, R., Akbari, O. S., 2020. Developing genetic tools to control ACP. Citrus Research Board. 11: 64–68.

Ellison, C.K., Niehuis, O., Gadau, J., 2008. Hybrid breakdown and mitochondrial dysfunction in hybrids of *Nasonia* parasitoid wasps. J. Evol. Biol. 21: 1844–1851.

Ferree, P.M., Fang, C., Mastrodimos, M., Hay, B.A., Amrhein, H., Akbari, O.S., 2015. Identification of genes uniquely expressed in the germ-line tissues of the jewel wasp *Nasonia vitripennis*. G3 5: 2647–2653.

Fuchs, H., Bachran, C., Flavell, D., 2013. Diving through membranes: molecular cunning to enforce the endosomal escape of antibody-targeted anti-tumor toxins. antibodies. Antibodies. 2: 209–235.

Heu, C.C., McCullough, F.M., Luan, J., Rasgon, J.L., 2020. CRISPR-Cas9-based genome editing in the silverleaf whitefly *(Bemisia tabaci)*. CRISPR J 3: 89–96.

Hunter, W.B., Gonzalez, M.T., Tomich, J., 2018. BAPC-assisted CRISPR/Cas9 system: Targeted delivery into adult ovaries for heritable germline gene editing (Arthropoda: Hemiptera). bioRxiv. 478743.

Kimura, K., 2001. Transposable element-mediated transgenesis in insects beyond *Drosophila*. J. Neurogenet. 15: 179–192.

Larracuente, A.M., Ferree, P.M., 2015. Simple method for fluorescence DNA in situ hybridization to squashed chromosomes. J. Vis. Exp. 6: 52288.

Leung, K., Zande, L., Beukeboom, L.W., 2019. Life-history traits of the Whiting polyploid line of the parasitoid *Nasonia vitripennis*. Entomol Exp. Appl. 167: 655–669

Li, M., Au, L.Y.C., Douglah, D., Chong, A., White, B.J., Ferree, P.M., Akbari, O.S., 2017a. Generation of heritable germline mutations in the jewel wasp *Nasonia vitripennis* using CRISPR/Cas9. Sci. Rep. 7: 901.

Li, M., Bui, M., Akbari, O.S., 2017b. Embryo microinjection and transplantation technique for *Nasonia vitripennis* genome manipulation. J. Vis. Exp. 130.

Lynch, J.A., Desplan, C., 2006. A method for parental RNA interference in the wasp *Nasonia vitripennis*. Nat. Protoc. 1: 486–494.

Macias, V.M., McKeand, S., Chaverra-Rodriguez, D., Hughes, G.L., Fazekas, A., Pujhari, S., Jasinskiene, N., James, A.A., Rasgon, J.L., 2020. Cas9-mediated gene-editing in the malaria mosquito by ReMOT Control. G3 10: 1353–1360.

Paolucci, S., Dalla Benetta, E., Salis, L., Doležel, D., van de Zande, L., Beukeboom, L.W., 2019. Latitudinal Variation in Circadian Rhythmicity in *Nasonia vitripennis*. Behav. Sci. 9.

Parker, E.D., Jr, Orzack, S.H., 1985. Genetic variation for the sex ratio in *Nasonia vitripennis*. Genetics 110: 93–105.

Raikhel, A.S., Dhadialla, T.S., 1992. Accumulation of yolk proteins in insect oocytes. Annual Review of Entomology. 37: 217–251.

Saunders, D.S., 1965. Larval diapause of maternal origin: Induction of diapause in *Nasonia vitripennis* (walk.) (Hymenoptera: Pteromalidae). J. Exp.Biol. 42, 495–508.

Sethuraman, N., O’brochta, D.A., 2005. The *Drosophila melanogaster cinnabar* gene is a cell autonomous genetic marker in *Aedes aegypti* (Diptera: Culicidae). Journal of Medical Entomology. 42: 716–718.

Sukthankar, P., Avila, L.A., Whitaker, S.K., Iwamoto, T., Morgenstern, A., Apostolidis, C., Liu, K., Hanzlik, R.P., Dadachova, E., Tomich, J.M., 2014. Branched amphiphilic peptide capsules: cellular uptake and retention of encapsulated solutes. Biochim. Biophys. Acta 1838:2296–2305.

Sukthankar, P., Gudlur, S., Avila, L.A., Whitaker, S.K., Katz, B.B., Hiromasa, Y., Gao, J., Thapa, P., Moore, D., Iwamoto, T., Chen, J., Tomich, J.M., 2013. Branched oligopeptides form nanocapsules with lipid vesicle characteristics. Langmuir 29: 14648–14654.

Valle, D., 1993. Vitellogenesis in insects and other groups--a review. Mem. Inst. Oswaldo Cruz 88: 1–26.

van de Zande, L., Verhulst, E.C., 2014. Genomic imprinting and maternal effect genes in haplodiploid sex determination. Sex Dev. 8: 74–82.

Verhulst, E.C., Beukeboom, L.W., van de Zande, L., 2010. Maternal control of haplodiploid sex determination in the wasp *Nasonia*. Science 328: 620–623.

Verhulst, E.C., Lynch, J.A., Bopp, D., Beukeboom, L.W., van de Zande, L., 2013. A new component of the Nasonia sex determining cascade is maternally silenced and regulates transformer expression. PLoS One 8: e63618.

Werren, J.H., 1983. Sex ratio evolution under local mate competition in a parasitic wasP. Evolution 37: 116–124.

Werren, J.H., Loehlin, D.W., 2009. The parasitoid wasp *Nasonia:* An emerging model system with haploid male genetics. Cold Spring Harb. Protoc. 134.

Werren, J. H. Sex ratio evolution under local mate competition in a parasitic wasp. Evolution 37: 116–124 (1983).

