## Supplementary information for "Germline mutagenesis of *Nasonia vitripennis* through ovarian delivery of CRISPR-Cas9 ribonucleoprotein"

**Supplementary Informations**

- Table S1
- Fig. S1- S3

**Table S1. Saponin injections**

Effect of different concentrations of saponin on survival and fecundity of injected females. Fecundity is calculated as the number of G0 offspring produced.

| Saponin concentration | Nº females injected | Survival Day 1 | Survival Day 2 | Survival Day 3 | Nº females laying eggs | Nº G0 adults | Diapause larvae |
|---|---|---|---|---|---|---|---|
| 0.0 ng/µl | 20 | 14 (70%) | 14 (70%) | 8 (40%) | 12 (60%) | 571 | 26 |
| 0.1 ng/µl | 76 | 44 (58%) | 34 (45%) | 34 (45%) | 16 (21%) | 451 | 191 |
| 0.5 ng/µl | 61 | 44 (72%) | 28 (46%) | 22 (36%) | 22 (36%) | 597 | 64 |
| 1.0 ng/µl | 82 | 58 (71%) | 51 (62%) | 17 (21%) | 38 (46%) | 792 | 233 |

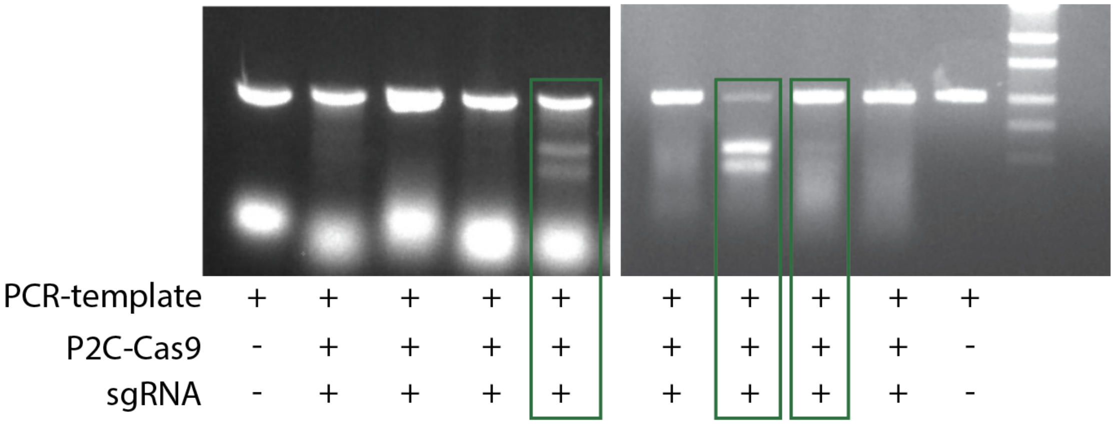


**Fig. S1 *In vitro* cleavage assay**

PCR of the targeted region of the *NV cinnabar* locus was used as a template for the *in vitro* cleavage reaction, pre-incubated with P2C-Cas9. The reaction is separated by agarose gel electrophoresis and different protein batches are present in different lanes. Green boxes highlight the protein batches that cleaved efficiently the target region *in vitro* and that were selected for further experiments.

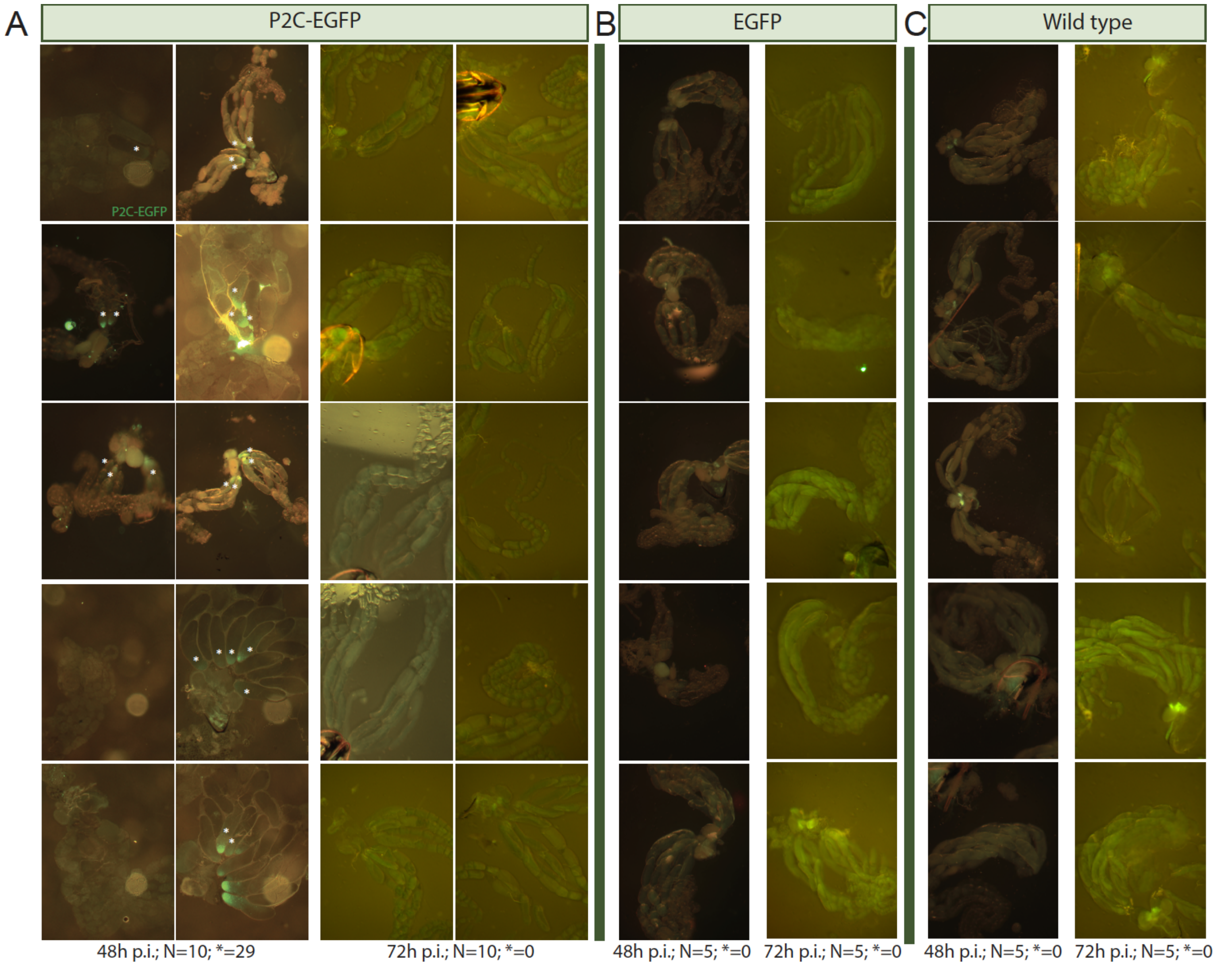


**Fig. S1 ovaries intake**

**A)** Delivery of EGFP by P2C into the ovaries of 48 and 72h post injection (p.i.). N represents the total number of ovaries and * the fluorescent-positive oocities. **B)** Delivery of EGFP without P2C peptides at 48 and 72h p.i. **C)** control non-injected ovaries.

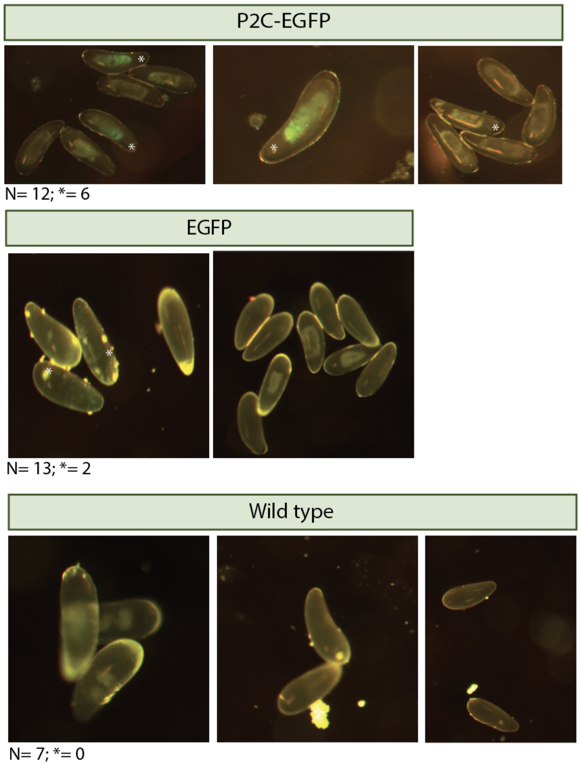


**Fig. S3 Embryos from 72h p.i. wasps**

Delivery of EGFP into embryos laid 72 h post injection (p.i.), respectively in P2C-EGFP, EGFP and non-injected treatments

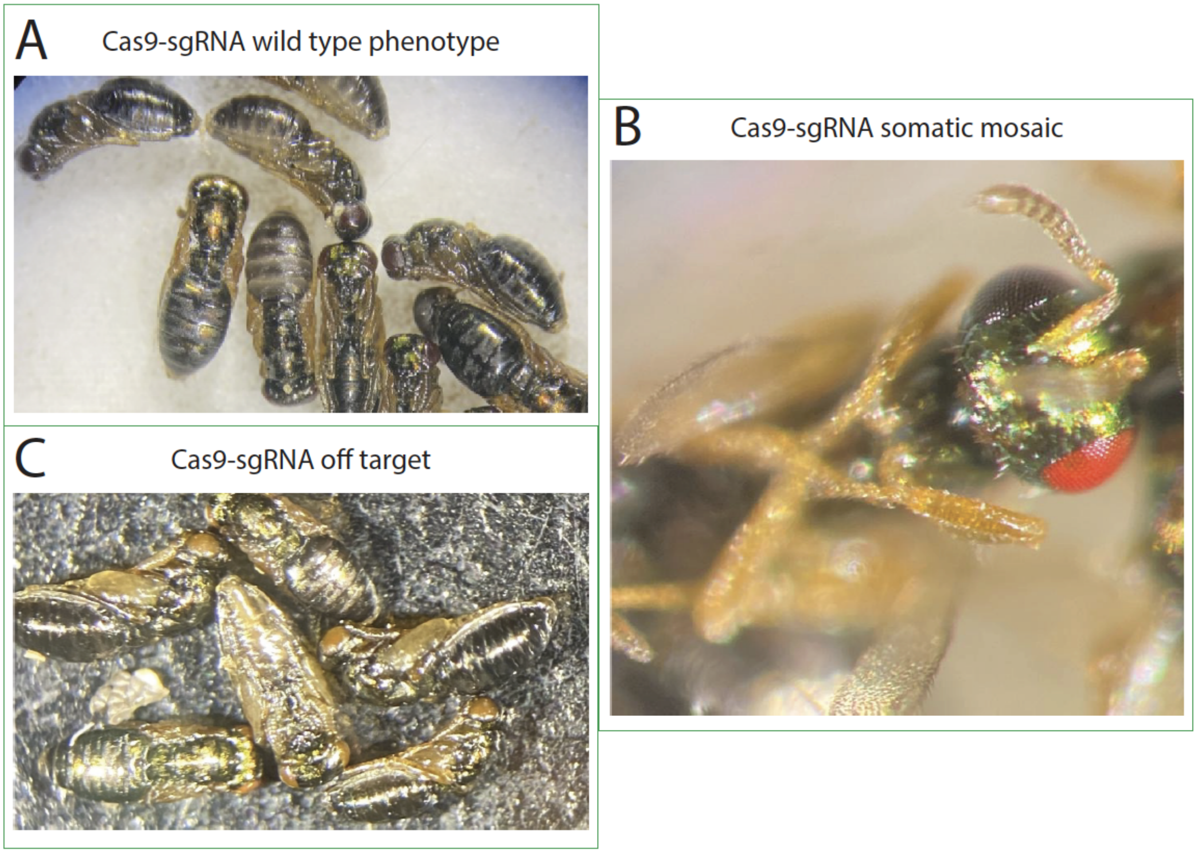


**Fig. S4 Cas9-sgRNA control injections**

**A)** Offspring with wild type phenotype produced by control injection Cas9-sgRNA. **B)** G0 mosaic mutant with one red eye and one wild type eye produced by control injection cas9-sgRNA **C)** Germline off target mutation appears as yellow/orange eyes after Cas9-sgRNA injection.
